# Global gene expression of human malaria parasite liver stages throughout intrahepatocytic development

**DOI:** 10.1101/2023.01.05.522945

**Authors:** Gigliola Zanghi, Hardik Patel, Nelly Camargo, Jenny L. Smith, Yeji Bae, Erika L. Flannery, Vorada Chuenchob, Matthew E. Fishbaugher, Sebastian A Mikolajczak, Wanlapa Roobsoong, Jetsumon Sattabongkot, Kiera Hayes, Ashley M. Vaughan, Stefan H. I. Kappe

**Author notes:** These authors contributed equally (Gigliola Zanghi, Hardik Patel, Nelly Camargo).

## Abstract

*Plasmodium falciparum* (*Pf*) is causing the greatest malaria burden, yet the liver stages (LS) of this most important parasite species have remained poorly studied. Here, we used a human liver-chimeric mouse model in combination with a novel fluorescent *Pf*NF54 parasite line (*Pf*NF54^*csp*^GFP) to isolate *Pf*LS-infected hepatocytes and generate transcriptomes that cover the major LS developmental phases in human hepatocytes. RNA-seq analysis of early *Pf* LS trophozoites two days after infection, revealed a central role of translational regulation in the transformation of the extracellular invasive sporozoite into intracellular LS. The developmental time course gene expression analysis indicated that fatty acid biosynthesis, isoprenoid biosynthesis and iron metabolism are sustaining LS development along with amino acid metabolism and biosynthesis. Countering oxidative stress appears to play an important role during intrahepatic LS development. Furthermore, we observed expression of the variant PfEMP1 antigen-encoding *var* genes, and we confirmed expression of PfEMP1 protein during LS development. Transcriptome comparison of the late *Pf* liver stage schizonts with *P. vivax* (*Pv*) late liver stages revealed highly conserved gene expression profiles among orthologous genes. A notable difference however was the expression of genes regulating sexual stage commitment. While *Pv* schizonts expressed markers of sexual commitment, the *Pf* LS parasites were not sexually committed and showed expression of gametocytogenesis repression factors. Our results provide the first comprehensive gene expression profile of the human malaria parasite *Pf* LS isolated during *in vivo* intrahepatocytic development. This data will inform biological studies and the search for effective intervention strategies that can prevent infection.

## INTRODUCTION

*Plasmodium falciparum* (*Pf*) is the causative agent of the most devastating form of human malaria, accountable for the vast majority of clinical cases and deaths [1]. Extensive malaria control efforts have significantly reduced disease morbidity and mortality in the last two decades. However, this decline has stagnated over the past seven years [2, 3] and more recently, has seen a rise to 241 million malaria cases and 627,000 deaths in 2021 [1]. While pointing towards the insufficiency of current efforts, the rise in disease incidence suggests the need of developing new interventions based on a better understanding of malaria parasite biology.

*Plasmodium* sporozoite (SPZ) forms are deposited into the skin of human hosts by infected *Anopheles* mosquito bite. The sporozoites then actively invade the bite-site-proximal blood capillaries and are carried via the bloodstream to the liver, where sporozoites cross the liver sinusoidal cell layer and invade hepatocytes. Sporozoites that have successfully invaded then transform into trophozoites and grow and replicate the parasite genome as a liver stage (LS) in a process called exo-erythrocytic schizogony [4]. During this phase the parasite undergoes massive cell size expansion and multiple rounds of genome and organellar replication while maturing into a late LS schizont that ultimately segments into tens of thousands of exo-erythrocytic merozoites. These red blood cell infectious forms egress from the infected hepatocytes, are released into the bloodstream and there initiate the symptomatic erythrocytic cycle of infection. The asymptomatic sporozoites and LS are considered the most promising target for malaria vaccine development. First, relatively low numbers of parasites are transmitted by mosquito bite and of these, only a fraction successfully infect the liver and undergo full LS development [5]. Second, the successful elimination of LS would prevent onset of symptomatic blood stage infection and onward transmission of parasites [6–8].

Numerous animal studies and clinical trials with live-attenuated parasites have demonstrated the importance of parasite developmental progression in the liver for eliciting broad, durable sterilizing immune protection [9–12]. For example, vaccination with fully infectious *Pf* sporozoites in combination with drugs that kill the parasite either in the liver or in the blood (Chemoprophylaxis vaccination, *Pf*SPZ-CVac) [12] confers superior protection in humans when compared to vaccination with radiation-attenuated *Pf*SPZ (RAS), which are unable to replicate in the liver. This difference in protective efficacy has been attributed to the notion that replication-competent *Pf*SPZ-CVac express a range of unknown LS antigens during their development in the liver, which are not expressed in replication-deficient PfSPZ-RAS.

Yet, despite the immunological and biological importance of *Pf*LS, gene expression of the parasite in the liver remains largely uncharacterized due to technical challenges. Notably, *Pf* SPZ have an almost exclusive tropism for primary human hepatocytes resulting in abnormal development in human hepatoma cell lines and low infection yields [13, 14]. Studies of *Pf* LS biology advanced with the use of *in vitro* primary hepatocyte culture systems [15] and the use of human liver-chimeric mouse models [16, 17]. The use of humanized mice has enabled *in vivo* studies of the pre-erythrocytic infection stages of *Plasmodium* species that infected humans. In particular, the fumarylacetoacetate hydrolase-deficient and immunocompromised FRGN mouse (Fah-/-, Rag2-/-, Il2rg-/-) transplanted with primary human hepatocytes (FRGN huHep mice) and transfused with red blood cells (huRBCs), are highly susceptible to infection with *P. falciparum* and *P. vivax* sporozoites and support full liver stage development and transition to blood stage infection [16, 18–20].

In the current study, we conducted a comprehensive transcriptomic analysis of *in vivo Pf* LS development. The fluorescent *Pf*NF54^*csp*^GFP line described herein enabled enrichment of *Pf* LS-infected primary human hepatocytes isolated from FRGN huHep mice. Our gene expression data show how the translational regulation machinery plays a key role in establishing LS infection and which genes and pathways are highly expressed during LS development. We further assessed how *Pf* and *Pv* LS schizonts are transcriptionally similar. Both species show similar gene expression profiles among orthologous genes, except the expression of genes involved in sexual stage commitment. Furthermore, *Pf* LS schizonts express several *var genes* [21], a class of clonally variant gene families (CVGs), that do not have orthologues in *Pv.* Our data sheds light on *Pf* LS gene expression serving as a basis for new avenues of vaccine and drug development.

## RESULTS

### *Pf* NF54^*csp*^GFP liver stages allow isolation of infected human hepatocytes

Gene expression analysis of *Pf* LS is encumbered by low hepatocyte infection rates and thus, low *Pf* LS mRNA representation compared to host mRNAs in total infected hepatocyte preparations as well as the lack of tools to isolate the infected cells. To overcome this issue, we created a fluorescent *Pf* parasite line that enables the isolation of parasite infected hepatocytes throughout LS development. We used the promoter region of the circumsporozoite protein gene *(CSP)* to drive strong expression of green fluorescent protein (GFP) and integrated this expression cassette into the dispensable *Pf47* locus to create the *Pf* NF54^*csp*^GFP parasite (Figure 1A). The *CSP* promotor was chosen because unlike CSP in rodent parasites [22, 23], *Pf* CSP is expressed throughout LS development [16, 19]. The *Pf* NF54^*csp*^GFP parasites were cloned and subsequently a single clonal population was extensively characterized at all the stages of the parasite life cycle (Supplementary Figure S1A and Table S1). The *Pf* NF54^*csp*^GFP parasites showed normal asexual blood stage growth and gametocyte maturation during blood stage culture. Furthermore, the infectivity of *Pf* NF54^*csp*^GFP parasites to mosquitoes was comparable to WT *Pf* NF54 parasites as evident from the oocyst counts in mosquito midguts and the enumeration of salivary glands sporozoites (Supplementary Figure S1B). Since the expression of CSP is initiated during oocyst development [24], we characterized CSP promoter-driven expression of GFP in the mosquito stages over time. By Western blot analysis the expression of GFP coincided with the endogenous expression of CSP (Supplementary Figure S1C). Live fluorescence microscopy further confirmed the expression of GFP in oocysts and salivary gland sporozoites (Figure 1B). Next, we assessed the competence of this parasite line to successfully complete the developmental cycle in the mammalian host. *Pf* NF54^*csp*^GFP sporozoites were by intravenously injected into FRGN huHep mice that had also been repopulated with huRBCs on day 6 and 7 post infection (PI). The *Pf* NF54^*csp*^GFP parasites were able to complete LS development and egress from the liver on day 7 PI. These transitioned parasites then grew normally in *in vitro* blood stage culture (Figure 1C).

**Figure 1:**
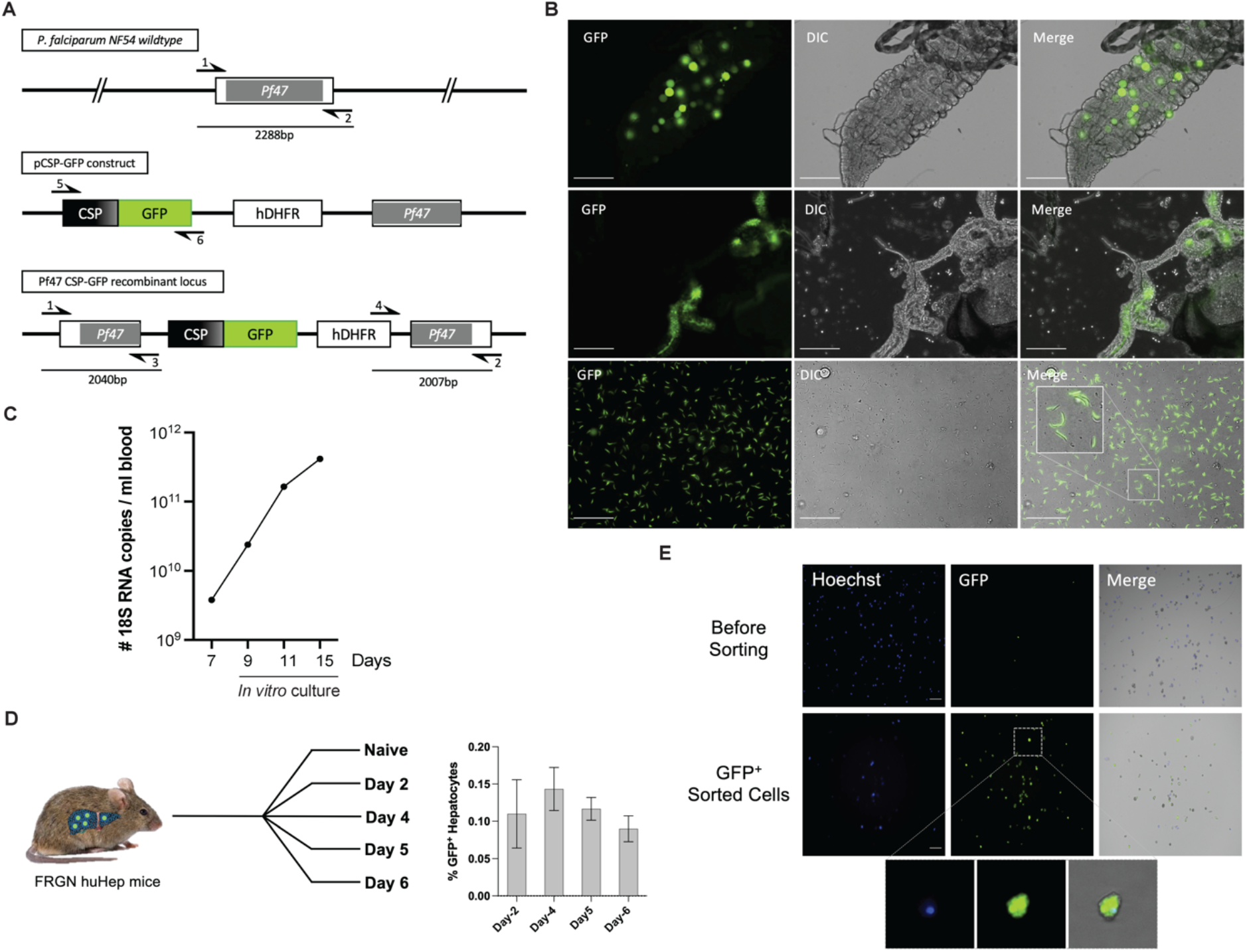
*The Pf NF54^csp^GFP* parasite enables the isolation of *Pf* infected primary human hepatocytes harvested from FRGN huHep mice. **(A)** Strategy to generate *Pf* NF54^CSP^GFP parasite line. Black arrows indicate PCR primer binding location that is used for the diagnostic PCR analysis (Supplementary Fig.1A). **(B)** Live imagining of *Pf* NF54^CSP^GFP in the mosquito stages to assess GFP expression, top panel Day 11 midguts, middle panel D14 salivary glands, lower panel D15 dissected sporozoites. Scale bar 50 μm. **(C)** *Pf* 18S rRNA measured at different time point by qRT-PCR after the blood was collected from FRGN huHep mouse infected with *Pf* NF54^CSP^GFP on day 7 PI and later cultured *in vitro.* **(D)** Experimental design. FACS Isolation of GFP^+^ *Pf* infected primary hepatocytes (*Pf*GFP^+^) from FRGN huHep mice on Day 2, 4, 5, or 6 post infection. Histogram shows percentage of *Pf* GFP^+^ hepatocytes at each time point (n=3) (Refer to Supplementary Fig. 1B). **(E)** Live imaging of GFP+ hepatocytes (green) before and after sorting. Nuclei stained by Hoechst dye (Blue).

For the isolation of *Pf* liver stages, female FRGN huHep mice, showing >70% primary human hepatocyte repopulation in their livers, were intravenously injected with 2.5 - 3 million *Pf* NF54^*csp*^GFP sporozoites per mouse. The mice were euthanized on days 2, 4, 5 or 6 PI and total primary hepatocytes were harvested by perfusing and digesting the livers [25]. To isolate a population of GFP^+^ *Pf* infected hepatocytes (*Pf*GFP^+^) and to minimize the contamination with uninfected cells that inherently possess high levels of autofluorescence, we applied a stringent gating strategy during FACS (Fluorescence activated cell sorting) (Supplementary Figure S2). We isolated *Pf*GFP^+^ hepatocytes from three biological replicates per time point with the average infection rate of 0.1 – 0.15% (Figure 1D) as per the FACS data analysis (Supp. Figure S2).

We confirmed the isolation and purity of *Pf*GFP^+^ infected hepatocytes from each batch by live fluorescent microscopy (Figure 1E). In total, we isolated between 2,500 – 3,000 *Pf*GFP^+^ and *PfGFP^-^* cells for each biological replicate and time point. The sorted cells were collected directly into lysis buffer, followed by RNA extraction, RNA-seq library preparation and next generation sequencing (NGS).

### The *P. falciparum* liver stage parasite transcriptome

We sequenced a total of 23 *Pf*GFP^+^ RNA-seq samples with biological and technical replicates. The 23 FASTQs were concatenated in 3 main biological replicates for each time point. We also generated a *Pf*NF54^*csp*^GFP sporozoite (*Pf*SPZ) transcriptome that was used along with four previously generated *Pf* sporozoite transcriptome data sets [26] as reference to comparatively analyze gene expression during *Pf* LS development.

All sample sequences were aligned to the *H. sapiens, M. musculus* and *P. falciparum* genomes. To reduce batch effects, we performed Combat-seq correction, which generates corrected gene counts retaining all sources of latent biological variation [27]. The average normalized expression of genes for each of the genomes (*Pf* or *Human*) showed that at the early time point of infection (day 2), human gene expression products were much more abundantly detected than those for *Pf* (Figure 2A). Representation of the *Pf* genome increased over time as the LS growth progresses in the liver reaching the highest value for late LS forms (Day 6). This contrasts with host coding reads, which remained relatively constant throughout parasite development (Figure 2A and Supplementary Table S2). This increase of the *Plasmodium* transcriptome representation over time is consistent with LS growth, genome replication and parasite expansion during exo-erythrocytic schizogony [28]. Using a cutoff of >= 1 CPM in the three biological replicates, we detected over 4000 expressed *Pf* genes for each time point (Figure 2B and Supplementary Table S3).

**Figure 2:**
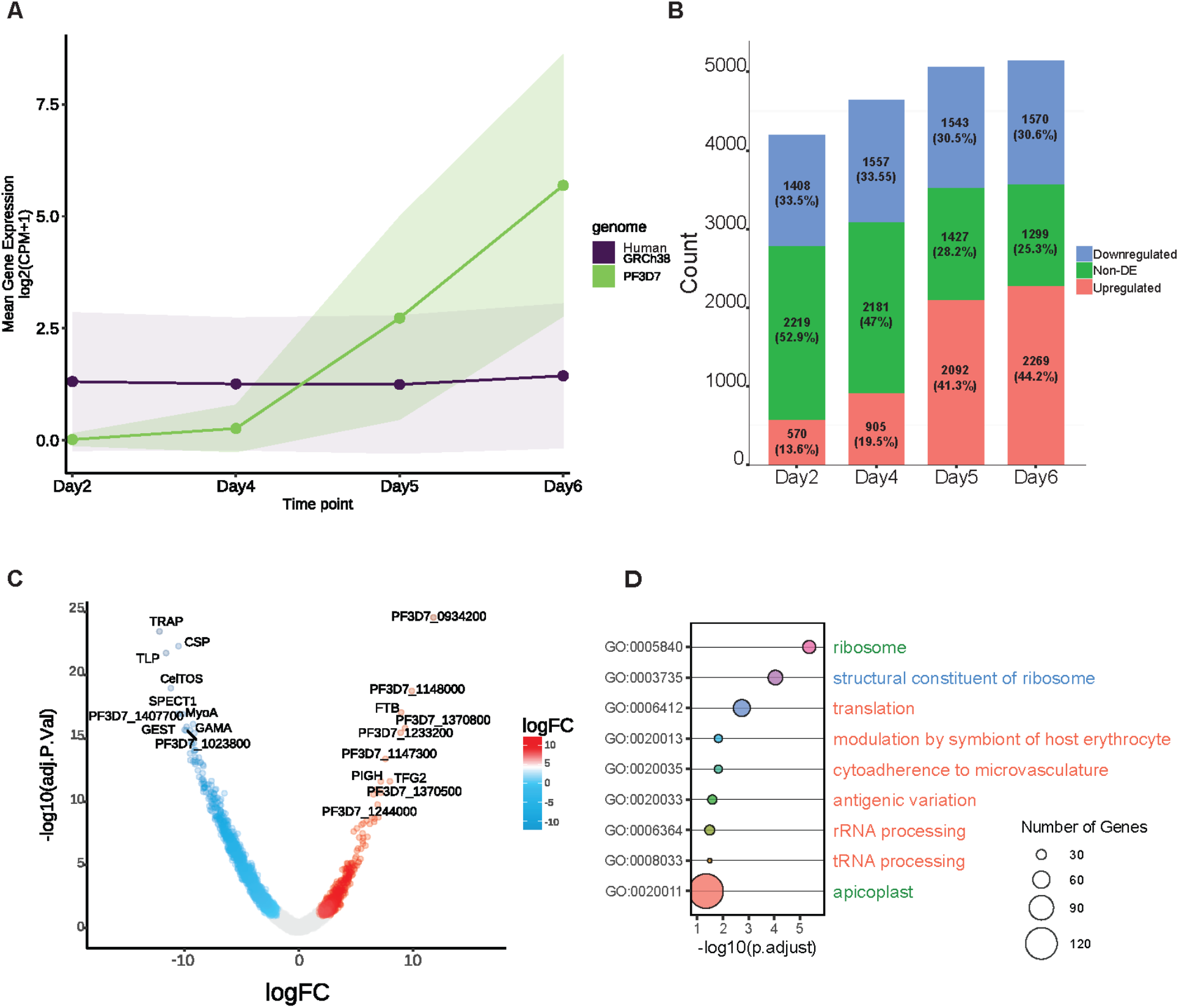
*P. falciparum* gene detected and early time point analysis. **A)** Ribbon graph showing the number of mean gene expression log2(CPM +1) aligning against *H. sapiens* (purple) and *P. falciparum* (green) genomes over time. **B)** Total number of genes detected at each time point and number of DEGs in reference to Sporozoites, genes considered expressed have >= 1 CPM in all 3 *Pf* biological. **C)** Volcano plot showing DEGs between Sporozoites (Blue) and *Pf* LS parasites at Day 2 PI (Red). **D)** Bubble plot showing GOTerm analysis of the upregulated genes at Day 2 PI. The size of the circles displayed is positively correlated with the number of genes involved in each pathway. Threshold in -log10 p-value.

Differential expression analysis comparison with sporozoites identified a total of 1978 differentially expressed genes (DEGs) at day 2 that increase over time, reaching a total of 3839 at day 6 (Figure 2B). During early (Day 2) and mid (Day 4) LS development most of these DEGs were lower in transcript abundance (~33%), with ~13 – 19% of transcripts increasing in abundance, whereas for the late LS (Day 5 and 6) transcript abundance increased to ~41 – 44% (Fig 2B) over sporozoites.

Invasion of hepatocytes by the motile sporozoite stage and the subsequent formation of the intracellular LS trophozoite stage constitutes a major transition point in the parasite life cycle. Transformation of the intracellular post invasion *Pf* sporozoite into a spherical trophozoite stage is completed approximately 2 days after infection and is characterized by a dramatic cellular remodeling process [29]. To explore the molecular mechanisms underlying this transformation, we performed differential gene expression analysis of the *Pf* LS Day 2 and the sporozoites stage. Among the 1978 DEGs identified at Day 2 PI, transcript abundance decreased for 1408 genes and increased for 570 genes compared to the sporozoite stage. Gene ontology enrichment analysis (Go Term) of genes for which transcript abundance decreased in LS trophozoites compared to sporozoites, revealed pathways involved in sporozoite motility (GO:0048870; GO:0071976) and cell invasion (GO:0044409; GO:0005515) (Supplementary Fig S3 and Supplementary Table S5). These included transcripts encoding sporozoite motility/cell traversal and invasion-related proteins, such as MyoA, CSP, CelTOS, TLP, GAMA and TRAP (Figure 2C and Supplementary Table S4). Genes for which transcript abundance increased in LS trophozoites compared to sporozoites, showed enrichment in pathways with a FDR ≤ 0.5 such as translational regulation (GO:0006412) and ribosome biogenesis (GO:0005840, GO:0003735) (Figure 2D). Interestingly, another pathway involved in the modulation of the host cells was upregulated at in LS trophozoites (GO:0020013). The genes enriched in this pathway are two classes of the CVGs, 14 members of the *rifin* genes and 1 member of the *stevor genes.* Furthermore, thirty of the upregulated gene products at day 2 were apicoplast targeted (GO:0020011) (Supplementary Table S5). These genes were further analyzed by metabolic pathway enrichment analysis, using KEGG and MetaCyc databases. This showed a preponderance of gene products mediating fatty acids biosynthesis and isoprenoid biosynthesis. Although the central metabolic role of apicoplast biosynthetic pathways was shown to be important during rodent malaria parasite LS development [30–32], their functional role as yet to be ascertained in *Pf* LS.

We next performed time course gene expression analysis to assess gene expression associated with different LS development stages. We excluded the day 2 LS time point from the time course analysis because this time point had relatively low parasite RNAseq reads when compared to the later LS time points. The mid-LS time-point (day 4) and the two late-LS time-points (days 5 and 6) were analyzed by time course cluster analysis using sporozoites as a reference. Clustering analysis of the z-score for all differentially expressed genes identified nine clusters (Figure 3A; Supplementary Table S6). Gene ontology (GO) analysis to identify biological process, found that ~25% of the genes expressed during LS development are not annotated, resulting in low statistical significance in the GO analysis. To increase statistical power, with more genes per cluster analyzed, we grouped the co-expression clusters by expression trend, reducing the number to five major cluster groups (Figure 3B and Supplementary Table S7). The first cluster group (Cluster 1 and 2) included genes for which transcript abundance was low throughout LS infection when compared to sporozoites such as pathways mediating cell motility and adhesion (GO:0071976, GO:0048870 and GO:0098609) (Figure 3C). The second cluster group (clusters 3 and 8) included genes that are expressed during LS but at lower levels in comparison to sporozoites. In this cluster group, the pathways were enriched for genes that play a role in entry into the host (GO:0044409), regulation of transcription (GO:0006355) and lipid metabolic process (GO:0006629) (Figure 3C). The third cluster group (clusters 4 and 6) included genes that are expressed in sporozoites but for which transcript abundance increased significantly (with a log2 FC ≥ 2) during LS development. This group included genes that *Plasmodium* expresses throughout the sporozoites and LS stages. The increase of transcript abundance for these genes during LS development can be explained by increased LS biomass with time. Indeed, the pathways enriched encoded key biological process such as RNA-binding (GO:0003723), ribosomes biogenesis (GO:0022625, GO:0005840, GO:0003735 and GO:00022627) and translation (GO:0006412, GO:0005852 and GO:0002181).

**Figure 3:**
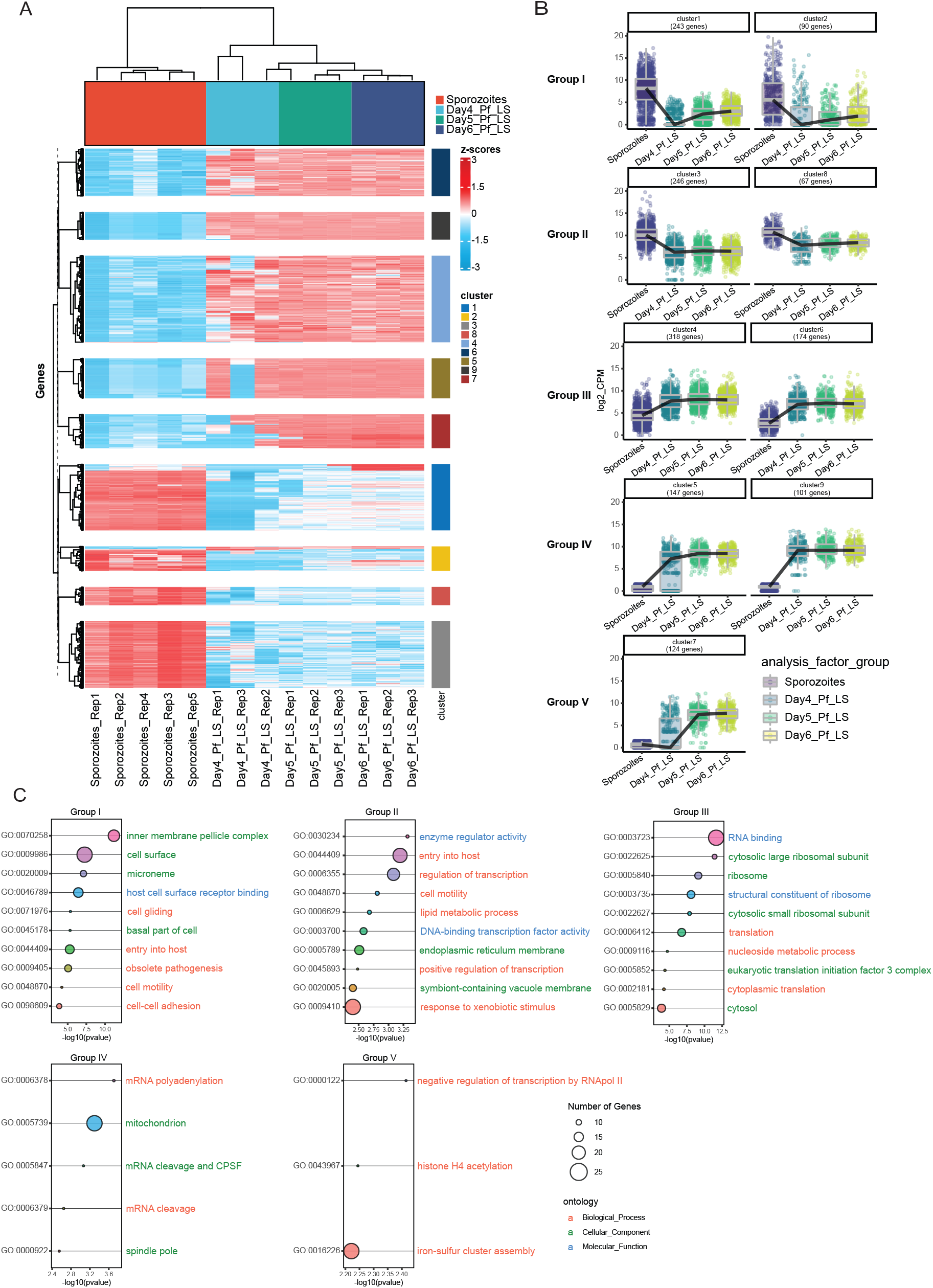
Time course cluster analysis of the *P. falciparum* LS transcriptome. **A)** Heatmap showing time course cluster analysis of *Pf* sporozoites (Red), Day 4 (Blue), Day 5 (Green) and Day 6 (Blue), gene expression values are shown as z-scores. **B)** Boxplot showing the expression trend of the nine cluster identified in panel A, the expression was further grouped in 5 main profiles. **C**) Bubble plot showing GOTerm analysis of the upregulated genes in the 5 expression profiles. The size of the circles displayed is positively correlated with the number of genes involved in each pathway. Threshold in - log10 p-value.

The fourth cluster group (Clusters 5 and 9) included the genes highly expressed during LS. The genes in this cluster are largely associated with mitochondria (GO:0005739). Metabolic pathway enrichment analysis of these genes identified pathways associated with amino acid metabolism. Furthermore, we observe upregulation of pathways involved in the response to oxidative stress (GO:0006979), indicating a parasite response to regulate redox homeostasis.

Cluster 7 is the sole member of the fifth cluster group characterized by genes upregulated during the mature stages of the LS. Most of the genes identified fall in the iron-sulfur cluster assembly (GO:0016226), showing how the iron metabolism could be essential during LS development. These results summarize the core LS transcriptome of *Pf* during an *in vivo* infection.

### Conservation of LS-specific gene expression among *P. falciparum* and *P. vivax*

To identify similarities in LS gene expression between the two most common *Plasmodium* species which infect humans, we compared the transcriptomes of late LS schizonts of *Pf* and *Pv.* Since the two species show differences in the duration of LS maturation, namely 6½ Days for *Pf and* 9 *Days* for *Pv* [16, 18], we selected *Pf* Day 6 and *Pv* at Day 8 as time points corresponding to late LS schizonts, as confirmed by the expression of MSP1 (data not shown).

To generate a late *Pv* LS transcriptome we could not utilize a fluorescent parasite that enables enrichment of LS-infected hepatocytes. Thus, the approach to generate the *Pv* LS transcriptome consisted of extraction of total RNA from 3 biological replicates of *Pv* infected FRGN huHep mice at Day 8 post infection, followed by enrichment for *Pv* mRNA by magnetic pull-down using custom made baits tiling the *P. vivax* P01 genome [33, 34]. Selection by hybrid capture was followed by Illumina sequencing. All sample sequences were aligned to the *H. sapiens, M. musculus* and *P. vivax* PO1 genomes as described above for the *Pf* dataset.

We first assessed the relatedness of *Pf* and *Pv* coding genomes. We identified that 78.4% or 4485 *Pf* genes have orthologues in *Pv.* We then considered expressed genes those with at least ≥1 TPM in all three biological replicates for each of the two species (Figure 4A, Supplementary Tables S9 and S10). We observed a substantial overlap of gene expression, with the majority of the orthologues (76%) expressed in late LS of both species. There were 334 genes exclusive to *Pv* LS, corresponding to the 8% of the total ortholog number. In contrast, there were 745 genes assigned as specific to the *Pf* coding genome (17%). Among the genes concordant in expression and gene ranking between the parasite species we found gene products that play an important role during LS development such as the liver specific protein 2 (LISP2) (14500 TPMs in *Pf* and 4904 in *Pv*) (Supplementary Tables S9 and S10). We further identified transcripts that show maturity of the LS schizonts MSP1 (3911 TPMs in *Pf* and 100 in *Pv*). GO term enrichment analysis for the genes co-expressed in the two species showed enrichment in pathways essential for parasite development, such mitochondria-targeted gene products (GO:005739) and protein transport associated gene products (GO:006886 and GO:0015031) (Supplementary Figure S4 and Table S11). In the gene products involved in the export pathway we identified transcripts of the parasitophorous vacuole (PVM) transporter EXP2 (306 TPMs for *Pf* and 234 TPM for *Pv*). EXP2 is a protein with dual function, one for nutrient transport and another for export of proteins across the PVM [35, 36].

**Figure 4:**
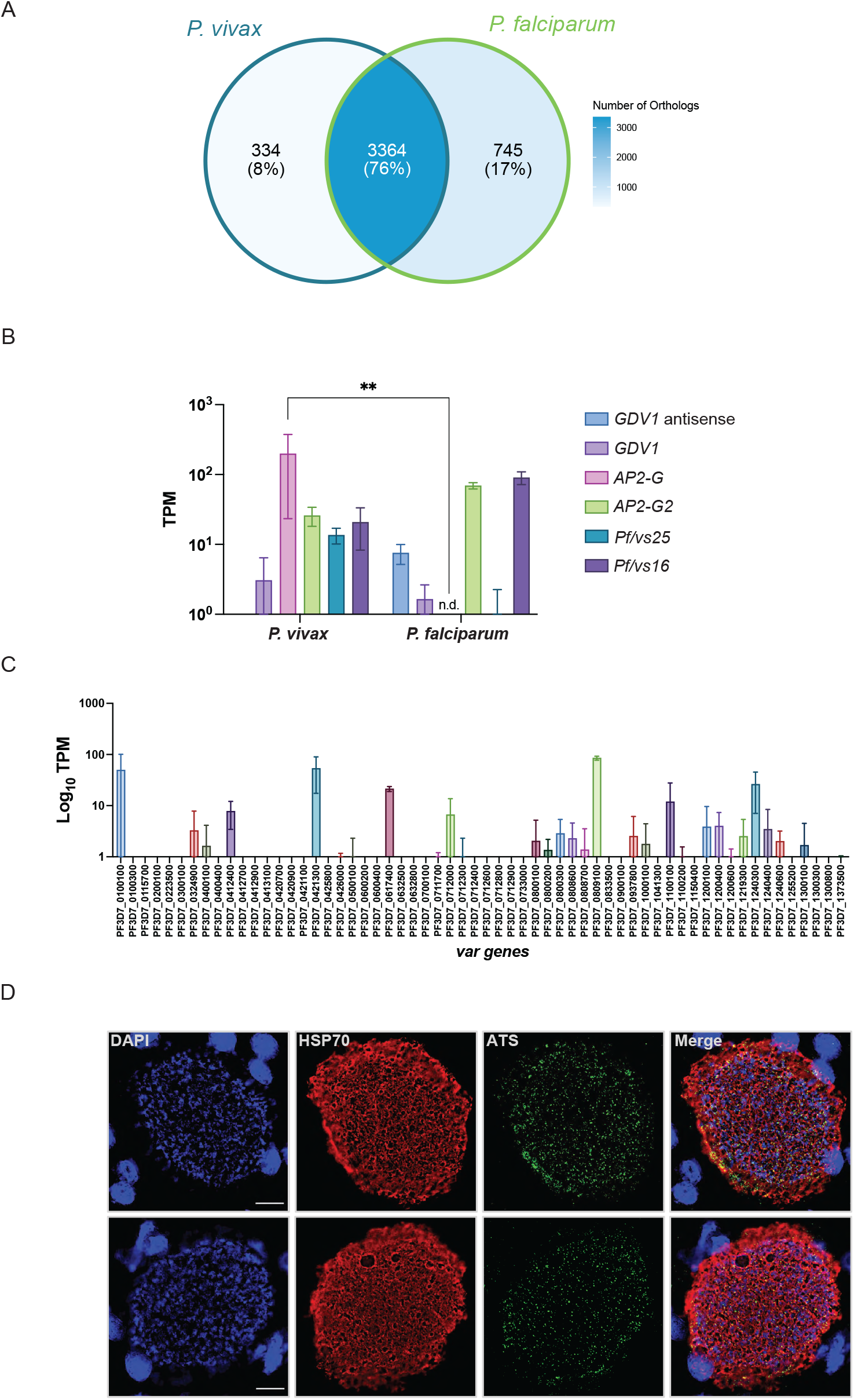
Similarities and differences of *Pf* and *Pv* LS transcriptome, and PfEMP1 expression in *P. falciparum* mature schizonts. **A)** Venn diagram of orthologues expressed genes with >= 1 TPM in all 3 biological replicates (*Pf* or *Pv*). Overlapping section identifies genes detected in the *P. falciparum* and *P. vivax* transcriptome. **(B)** TPM values of a selection of gametocytes genes in the *Pf* and *Pv* datasets. **C)** TPM values of the var genes detected in the *P. falciparum* transcriptome. **D)** Liver stage parasites. Both panels, shows *Pf* liver stage schizonts from FRGN huHep mice immunostained with DAPI (blue), anti-PfHSP70 (red) and anti-PfATS at day 7 post-infection. The scale bar corresponds to 10 μm.

We further assessed whether *Pf* LS schizonts are sexually committed before egressing from the liver, as previously shown at the transcriptomic level for the *Pv* LS schizonts [37–40]. Thus, we first evaluated detection of gametocytes commitment genes in the *Pv* Day 8 LS schizonts. We detected the expression of *Pvs16* and *Pvs25* along with the *AP2-G* transcription factor that regulates sexual commitment [41–43] (Figure 4B). In contrast, we did not find the expression of *AP2-G* in the *Pf* dataset. Interestingly, we found that *Pf* Day 6 LS schizont expressed the antisense transcripts of *GDV1* (Figure 4B). The *GDV1* antisense transcripts controls the expression of the *GDV1* gene, that operates upstream of *AP2-G,* acting as a master regulator that induces sexual differentiation [44]. These results together suggest that *Pf* does not commit to sexual stage development in the liver. Among the genes that are expressed in *Pf* LS that do not have orthologues in *P. vivax,* we found expression of multiple members of the *var* gene family (Figure 4C). The *var* genes encode the erythrocyte-membrane protein-1 (PfEMP1) adhesin family, which mediates both antigenic variation and cytoadherence of infected erythrocytes to the microvasculature [45]. Although *var* gene expression is commonly associated with asexual blood stages, there is evidence that *var* gene members can be transcribed in gametocytes, ookinetes, oocysts, and sporozoites [26, 46–48].

In our data we found transcripts of multiple *var* expressed at the same time. Notably, among the expressed *var* (Figure 4C), we found consistent expression of 10 *var* genes in all 3 biological replicates, belonging to the the UpsB and UpsC types [49]. The PF3D7_0809100 *var* gene, previously shown to be expressed in sporozoites [26], appeared to be the most highly expressed in late LS schizogony. We further investigated PfEMP1 protein expression in *Pf* liver stage schizonts. We used a pan antibody recognizing the PfEMP1 semi-conserved intracellular region ATS (red), to stain day 7 LS. A punctate staining pattern was observed (Figure 4D), confirming PfEMP1 expression in *Pf* liver schizonts. All together these findings suggest that resetting of *var* gene expression initiates in the mosquito stages and concludes at the end of the liver stage development, possibly explaining why a broad repertoire of *var* genes in the first generations of blood-stage parasites is observed in malaria-naive human volunteers infected with *Pf* sporozoites [50, 51].

## DISCUSSION

LS are critical targets for vaccine and drug development to prevent the onset of symptomatic blood stage infection and onward parasite transmission of the parasite by the mosquito vector. Transcriptome analyses conducted on *in vitro* non_*Pf* LS parasites have been instrumental in revealing gene expression, LS-specific biological processes [23, 40, 52–54] and providing important insight, including hypnozoite activation markers [37, 55], and comparative gene expression analysis with other stages, even at a single-cell resolution [40, 56]. Yet, the gene expression profiles for the LS of the most medically important *Pf* parasite that grow within infected human hepatocytes had not yet been reported. Here we provide the *in vivo* analysis of *Pf* LS gene expression throughout their six-day intra-hepatocytic development by performing transcriptome analysis. To date, the analysis of *Pf* LS-infected hepatocytes has been extremely challenging, mostly due to low infection rates, the almost exclusive trophism for primary human hepatocytes and the inability to effectively isolate infected cells. Furthermore, although several *Pf* fluorescent lines are available and have greatly served the cellular and molecular characterization of the different stages of the parasite, their fluorescence did not enable genome-wide transcriptomic analysis of the *Pf* intrahepatic stages. *The Pf* NF54^*csp*^GFP parasite line described herein allowed us to isolate pure populations of *Pf* LS-infected human hepatocytes from FRG huHep mice at different time points of infection in the liver and also demonstrated that the *Pf CSP* promoter remains active throughout LS development.

Our transcriptional analysis unraveled the genes and pathways driving *Pf* LS biology. Approximately 25% of genes identified as expressed in the LS transcriptome had no available GO term annotations. Illustrating the importance of further investigating the unknown function of a ¼ of the *Pf* genome during LS development. Notwithstanding, we identified the core pathways the parasite uses to develop within the host hepatocyte.

At an early time point of infection, the parasite establishes infection of the host cell by transforming from the sporozoite stage to the LS trophozoite [4]. Transition from the vector to the host requires rapid translation of proteins needed for mammalian host infection. At Day 2 PI, we found downregulation of the genes associated with the translational repression machinery and upregulation of genes essential to support LS formation and replication [57, 58]. Furthermore, our data set provides gene expression patterns of LS genes that sustain the drastic metamorphosis the parasite undergoes to enter rapid replication. During LS development the central metabolic pathways are involved in fatty acid and isoprenoid biosynthesis that are enriched as soon as Day 2 PI [23, 59, 60]. These pathways along with the amino acid synthesis and metabolism are then drastically upregulated at later time points of the LS development. Other important gene products, enabling parasite growth within the host cell, are involved in redox homeostasis, which the parasite likely utilizes to counteract the oxidative stress generated in the infected host cell [23, 61].

We further analyzed the differences in gene expression between *Pf* and *Pv* LS schizonts. Our data are in agreement with previous findings and observations that *Pv* produces gametocytes as soon as egressing from the liver resulting in rapid transmission [37–40, 62]. The *Pv* dataset supports the hypothesis of commitment to gametocytogenesis in *Pv* LS schizonts, leading to formation of sexually committed merozoites. This property is not conserved in the *Pf* schizonts, which appear to not be sexually committed. This is in accordance with the current understanding of *Pf* gametocytogenesis in infected individuals. Furthermore, in the *Pf* LS transcriptome we observed expression of the antisense *GDV1* gene [44]. It has been shown that this antisense transcript, inhibits the transcription of *GDV1* that acts as regulator of *AP2-G* which in turn regulates sexual commitment. These findings indicate that *Pf* LS schizonts have not initiated de-repression of *AP2-G* expression [44]. Thus, exo-erythrocytic *Pf* merozoites when egressing from the liver are not sexually committed.

Among the genes conserved in the *Pf* and *Pv* datasets, we found expression of EXP2. The expression of this small-molecule transporter is critical during LS development in rodent malaria parasites [63]. Recent rodent malaria studies show that EXP2, during LS, retains some function as part of the export machinery (PTEX) even if differently from the blood stage due to the absence of some translocon components [63]. However, the function of EXP2/PTEX is less clear during LS. Further analyses are required to understand the possible role of EXP2 in LS parasite protein export across the PVM.

Furthermore, we characterized the transcriptome of the *Pf var* gene family encoding antigenically variant PfEMP1 proteins, which are the major determinates of *Pf* malaria pathology and immune escape during blood stage replication [45]. It is becoming increasingly apparent that *Plasmodium* variant multigene families are not exclusively associated with blood-stage infection and may play additional roles across the life cycle. Indeed, we identified multiple members of the *var* gene family in the *Pf* LS transcriptome. Immune evasion relies on the antigenic variation depending on monoallelic expression of one *var* gene at any given time. So far it has not been formally shown where the *var* gene transcription resetting occurs. It has been speculated to take place either in the vector, where studies with human volunteers have shown that *var* gene repertoire is altered upon mosquito transmission, or during pre-erythrocytic stages [50].

Here we demonstrate *var* gene clonal deregulation and apparent monoallelic expression disruption during LS development. To validate the resetting model during LS development, future single cell profiling of the *var* gene repertoire in individual liver stages will have to be conducted. Due to the prevalence of post-transcriptional repression mechanisms in *Plasmodium*, it remained however unclear whether *var* transcriptional activity translates into protein expression. To assess if transcripts were translated into PfEMP1s we analyzed expression at the protein level and did observe PfEMP1 protein expression in fully mature LS schizonts. One possible role of the PfEMP1 expressed in LS schizonts, is based on the known adhesive functions of this protein family. PfEMP1s might be exported to the merosomal membrane to bind to the pulmonary endothelium expressing CD36. This hypothesis, might explain the high efficiency of merosomes arrest and merozoite release in the lung vasculature, as shown in rodent malaria parasites [64]. However, more in depth studies are needed to confirm role of PfEMP1 expression during LS infection.

In conclusion our work offers a comprehensive view of the *P. falciparum* LS transcriptome *in vivo* and our comparative analysis with the *P. vivax* late LS transcriptome pinpointed the common genes expressed during LS development in both species. These findings identified new cross-species candidates valuable for the development of new intervention strategies. Future studies will further advance our molecular understanding of this critical stage in the *Plasmodium* life cycle.

## ACKNOWLEDGMENTS

We thank the vivarium staff at Seattle Children’s Research Institute for their constant support during animal studies. In addition, we thank the insectary staff for rearing the mosquitoes for these studies. This work was funded by a NIH U01 AI155335 to S.H.I.K and a BMGF grant (OPP1137694) to S.H.I.K and S.A.M.

## AUTHOR CONTRIBUTIONS

Conceptualization, G.Z., H.P., E.L.F., S.A.M., S.H.I.K.; Methodology, G.Z., E.L.F., H.P.; Investigation, G.Z., H.P., N.C., J.L.S., Y.B., E.L.F., V.C., M.E.F, S.A.M., K.H., A.M.V., S.H.I.K.; Writing, G.Z., H.P., A.M.V., S.H.I.K.; Resources and Funding Acquisition, S.H.I.K. and S.A.M.

## DATA AVAILABILITY

GEO accession number pending.

## MATERIALS AND METHODS

### Creation of *Pf* parasite line expressing GFP under CSP promoter

The plasmid pEFGFP used to created 3D7HT-GFP was modified by replacing EF1α promoter with 1.2 kb DNA fragment from 5’ UTR immediately upstream to the start codon of *Pf* CSP gene (CSP promoter). Plasmid integrity was confirmed by DNA sequencing and used for transfection of *Pf* NF54.

*Pf* NF54 parasite culture was synchronized at ring stage with 5% sorbitol two days prior to transfection. On the following day trophozoites were selected by incubation in 0.7% gelatin solution. Ring stages were transfected by electroporation at 0.31 kV and 950 μF with a Bio-Rad Gene Pulser (BioRad, La Jolla, CA) as described previously (REFERENCE). Cultures were put under drug pressure starting at 6 hours post-transfection using 5nM WR99210 (Jacobus Pharmaceuticals). Integration was confirmed by PCR on parental population and clones obtained by limiting dilution as previously described using primers detailed in supplementary table 1.

### Mosquito Rearing and Sporozoite Production

*Anopheles stephensi* mosquitoes were reared and maintained following standard procedures outlined in Methods in Anopheles Research MR4. Mosquitoes were kept at 27°C and 75% humidity in temperature and humidity-controlled incubators on a 12 hours light/dark cycles within a secured ACL2 Facility. Cotton pads soaked in 8% dextrose and 0.05% PABA solution were placed daily on the top net of mosquito cages. *Pf* NF54 ^CSP^GFP asexual parasites were maintained by subculturing at 2% parasitemia in RPMI 1640 (25 mM HEPES, 2 mM L-glutamine media containing 10% human serum and 50 μM hypoxanthine and maintained at 37°C in an atmosphere of 5% CO2, 5% O2, and 90% N2.

Gametocyte cultures were set up at 1% parasitemia and 5% hematocrit, the asexual parent cultures had a parasitemia between 3-7%. The media of the gametocyte cultures were changed daily for 15 days while keeping the plates/flask on a slide warmer during changing the media to prevent dramatic temperature changes. Mature gametocytes cultures were spun at 800g for two minutes at 37°C in a temperature-controlled centrifuge.

Parasitized red blood cell pellet was re-suspended at 0.5% gametocytemia in 50:50 serum: blood mix and used for standard membrane feeding as described [65]. For *Pf* sporozoites production, 3-7 days old female mosquitoes were used for every infectious blood meal. Mosquito infections were evaluated on day 7 by checking oocyst prevalence and oocyst number in 10-12 dissected mosquito midguts. Sporozoite numbers were determined by dissecting and grinding salivary glands on days 15 post feed. These sporozoites were used for infecting FRGN huHep mice.

### Mice

FRG NOD huHep mice (female, >4 months of age) were purchased from Yecuris, Inc. and were housed and maintained in pathogen-free BSL2+ animal facility at the Center for Global Infectious Disease Research, Seattle Children’s Research Institute (SCRI). All animal procedures were performed as per the regulations of the SCRI’s Institutional Animal Care and Use Committee (IACUC). The animal procedures were approved by IACUC under 00480 protocol. Repopulation of human hepatocytes in FRGN huHep mice was confirmed by measuring human serum albumin levels, and only animals with human serum albumin levels >4 mg/mL corresponding to 70% repopulation of human hepatocytes were used as previously described [16]. Animals were cycled on 8 mg/L of NTBC once a month for 4 days to maintain hepatocyte chimerism. Mice were taken off from NTBC drug prior to and during experimentation.

### Analyzing the *Pf* NF54^CSP^GFP liver stage-to-blood stage transition in FRGN huHep mice by quantitative RT-PCR (qPCR)

FRGN huHep mice were intravenously (IV) infected with 1 million *Pf* NF54^CSP^GFP sporozoites. To support the parasites transition from liver stage-to-blood stage, mice were injected with 400 μl of human RBCs IV at 70% hematocrit on days 6 and 7 post infection. The blood was then collected by cardiac puncture after exsanguinating mice on day 7. Fifty microliters of blood were added to NucliSENS lysis buffer (bioM rieux, Marcy-l’ toile, France) and frozen immediately at −80°C and the rest of the blood was transferred to *in vitro* culture. Fifty microliters blood samples were collected from *in vitro* culture on day 9, 11 and 15 post infection in mice (i.e., day 2, 4 and 8 of *in vitro* culture) and were added to NucliSENS lysis buffer and frozen at −80°C. All the samples were processed and analyzed for presence of 18S rRNA as follow. The qRT-PCR reaction was performed using 35 μL SensiFAST™ Probe LoROX One-Step Kit (Bioline, Taunton, MA) and 15 μL of extracted eluate. Plasmodium 18S rRNA primers/probes (LCG BioSearch Technologies, Novato, CA) were as follows: Forward primer PanDDT1043F19 (0.2 μM): 5’-AAAGTTAAGGGAGTGAAGA-3’; Reverse primer PanDDT1197R22 (0.2 μM): 5’-AAGACTTTGATTTCTCATAAGG-3’; Probe (0.1 μM): 5’-[CAL Fluor Orange 560]-ACCGTCGTAATCTTAACCATAAACTA[T(Black Hole Quencher1)]GCCGACTAG-3’[Spacer C3]). Cycling conditions were RT (10 min) at 48°C, denaturation (2 min) at 95°C and 45 PCR cycles of 95°C (5 sec) and 50°C (35 sec).

### Isolation of *Pf* NF54^CSP^GFP LS infected primary human hepatocytes

The FRGN huHep mice were intravenously injected with 3 million *Pf* NF54^CSP^GFP SPZ per mouse. The primary hepatocytes were harvested by perfusing and digesting the livers on day 2, 4, 5 or 6 post sporozoites infection using modified protocol [25]. Briefly, the mice were deeply anesthetized with ketamine (100 mg/kg body weight)/Xylazine (10 mg/kg body weight) solution and the livers were perfused and digested with perfusion buffers I (0.5 mM EGTA in 1x DPBS without Ca^2+^ and Mg^2+^) and II (50 μg/ml liberase TL with 800 μM CaCl2 in 1x DPBS), respectively. The hepatocytes were dispersed in 1x DMEM complete medium and centrifuged twice at low speed (50 rpm) for 2 min at 10°C. The cell pellet was resuspended in the complete medium and the viability was tested using trypan blue staining. The final cell concentration was adjusted to 2 x 10^6^ / ml and further used for the FACS (Fluorescence activated cell sorting). Two thousand five hundred to three thousand GFP+ and GFP-cells were sorted directly in to the QIAzol lysis buffer.

### RNA-seq Library Preparation

For *P. falciparum* (*Pf*) RNA-seq libraries preparation, total RNA was extracted from sorted infected primary human hepatocytes using miRNeasy Micro Kit (QIAGEN) according to the manufacturer’s instructions, including on-column DNase digestion. Libraries were prepared using SMART-seq v4 Ultra Low Input (Clontech) and were sequenced on the Illumina NextSeq 500 as 75-bp pair-end reads. The resulting data were demultiplexed using bcl2fastq2 (Illumina) to obtain fastq files for the downstream analysis. A minimum of three biological replicates were analyzed; technical replicate libraries for each biological replicate were also sequenced. Additional raw sequence reads from *Pf* sporozoite RNA-seq samples (N=4) were retrieved from the Sequence Read Archive (PRJNA344838) [26].

For *P. vivax* (*Pv*) RNA-seq library preparation, total RNA was extracted from the FRGN huHep mice livers infected with 1 million *Pv* sporozoites (field strain) using TRIzol (Thermo Fisher) and purified using RNeasy Mini Kit (Qiagen) according to manufacturer’s instructions. A SureSelect XT custom oligo library was designed with Agilent (Design ID: S0782852) to enrich for *Pv* specific cDNA among the pool of human, mouse and parasite cDNA obtained from the RNA extraction from the humanized mouse liver. Total 85,000 probes of 120 bp size were tiled every 100 bp across the entire *Pv* Sal I genome. Sequences >30% similar to human sequences were excluded. Sequencing libraries were prepared according to the SureSelect XT RNA Target Enrichment for Illumina Multiplexed Sequencing protocol from Agilent (Ref: 5190-4393). Libraries were analyzed using a BioAnalyzer and were quantified using qPCR. Illumina libraries were sequenced on Mi-seq as 75-bp single-end reads.

### Data Analysis

Quality control of fastq files was performed using FastQC software; fastqs from paired biological and technical replicates of the liver-stage samples were concatenated to increase sequencing depth and coverage. *Pf* liver-stage and sporozoite samples’ sequencing reads were mapped to a reference genome containing *H. sapiens* (Ref, GRCh38, Ensembl gene annotations v106)*, M. musculus* (Ref, GRCm39, Ensembl gene annotations v106) and *P. falciparum* genome (Gardner et al., 2002, PlasmoDB, PlasmoDB-58_Pfalciparum3D7) with STAR 2.7.9. The alignment was completed with default parameters with the addition of “--twopassMode Basic” for high quality splice junction quantification and “--quantMode GeneCounts” to produce a gene count matrix. Gene level counts were normalized to counts per million (CPM) or transcripts per million (TPM). The *P. vivax* samples were processed identically, except with the reference genome containing *H. sapiens, M. musculus,* and the *P. vivax* genome (PlasmoDB-58_PvivaxP01).

### Differential Expression and Clustering analysis

All analyses were conducted in the R v4.1 statistical environment. The raw gene count matrix for *Pf* LS and sporozoite samples, including publicly available sporozoite data from PRJNA344838, underwent batch correction using ComBat-seq (sva v3.42.0) with default parameters. The adjusted counts were used in differential gene expression analysis with Limma voom v3.50.3; genes with absolute log2 fold change > 1 and false discovery rate (FDR) < 0.05 were retained. Batch corrected gene counts were trimmed mean of M-values (TMM) normalized and converted to log2 scale prior to differential expression analysis, unsupervised hierarchical clustering, and time-course regression analysis [66]. Changes in *Pf* LS gene expression over time compared to sporozoites was conducted using maSigPro v1.66.0 with a polynomial degree of 2 (quadratic). Comparison of *Pf* and *Pv* samples were carried out with TPM normalized gene-counts (unadjusted). For each species independently, genes were selected if expressed ≥1 TPM in all 3 biological replicates; the expressed genes were converted into *Pf* orthologs using PlasmoDB v58. The orthologs from *Pf* and *Pv* were overlapped and visualized using ggVennDiagram v1.2.2. Gene ontology analyses were performed using the GO enrichment tool for Biological Processes, Cellular Component and Molecular Function using ClusterProfiler v4.2.2. Available GO terms for *Pf* were downloaded from PlasmoDB v58.

### Live Imaging and Immunofluorescence assay

#### Live Imaging

Live images were captured using Keyence BZ-X700 fluorescence microscope and Fiji software was used for the image analysis.

#### Immunofluorescence

Livers were harvested from *Pf* infected FRGN huHep mice on day 7 post infection, fixed in 4% (vol/vol) paraformaldehyde (PFA, Alfa Aesar) in 1x PBS. Fifty-micron sections of the liver were blocked in normal goat serum diluted 1:500 for 2 h at 37 °C, washed twice in PBS and incubated overnight with anti-PfHSP70 antibody (1:1000) and anti-PfATS antibody (1:500), washed twice in PBS, then incubated with secondary antibodies and DAPI at 37 °C for 2 hrs. All antibodies were diluted in PBS containing 1% BSA and 0.2% Triton X-100. All sections were washed twice in PBS before being mounted in anti-fading medium and stored at 4 °C before analysis. Images were captured using the GE DeltaVision Elite optical/digital-sectioning fluorescence microscope, and Fiji software was used for the image analysis.

## SUPPLEMENTARY INFORMATION

**Supplemental_Table_S1_Primer sequences**

**Supplementary_Table_S2_PfLSTranscriptome_counts**

**Supplementary_Table_S3_PfLSTranscriptome_CPM**

**Supplementary Table S4_Differentialgeneepression_PfSPZ/Day2**

**Supplementary Table S5 ClusterProfiler**

**Supplementary_Table_S6_timecourse_ ClusterProfiler**

**Supplementary_Table_S7_timecourse_ ClusterProfiler _combined**

**Supplementary_Table_S8_timecourse_ ClusterProfiler _GoTerm**

**Supplementary Table S9_Pvivax_day8_Orthologues**

**Supplementary Table S10_Pfalciparum_day6_Orthologues**

**Supplementary_Table_S11_orthologs_go_ClusterProfiler_AllOntology**

## SUPPLEMENTARY FIGURE LEGEND

**Supplementary Figure S1:**
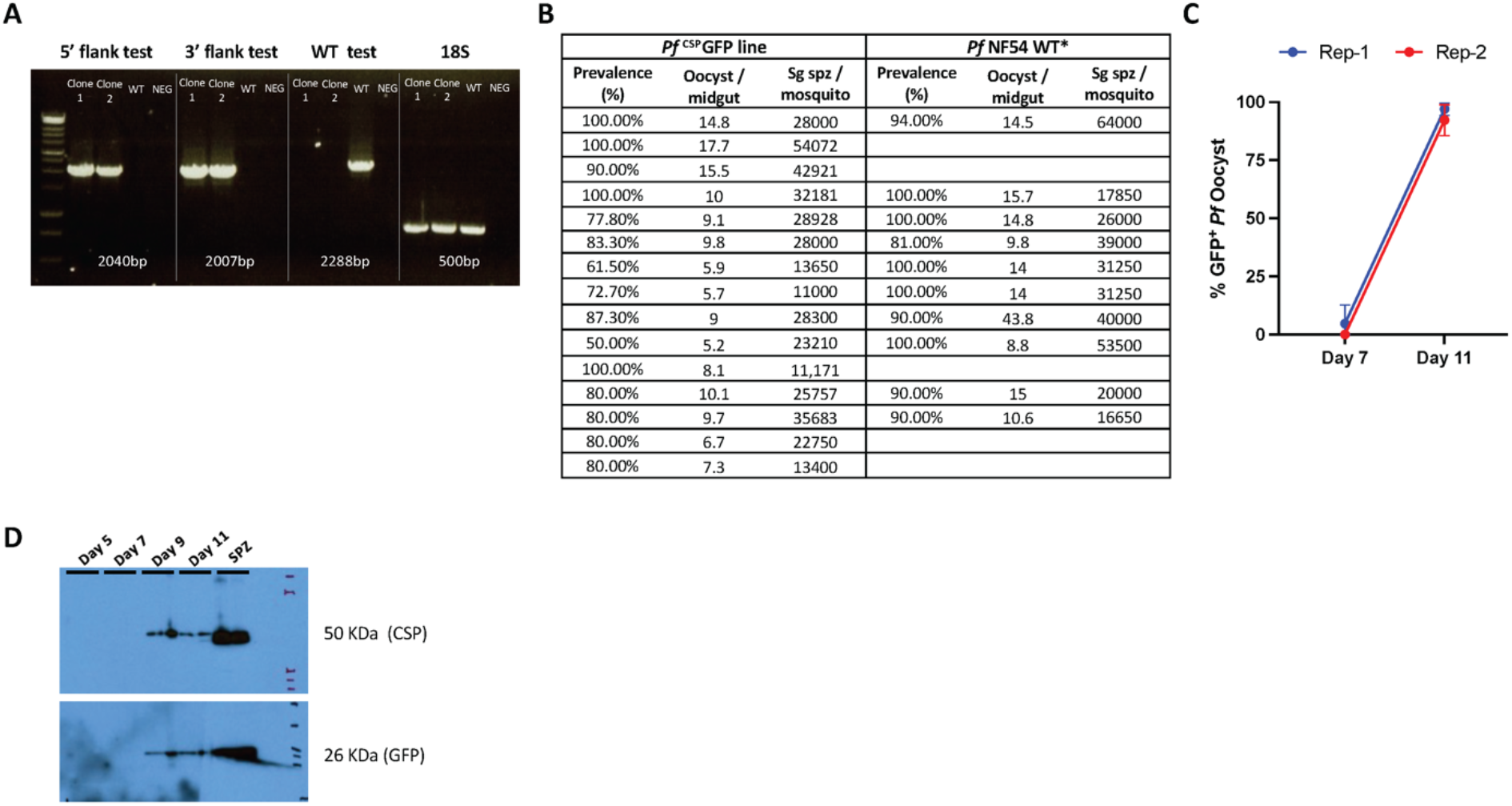
*P. falciparum* ^CSP^GFP *parasites display normal characteristics throughout life cycle.* **A)** Gel shows PCR results with primer sets to amplify recombinant DNA template of pCSP-GFP construct and show 5’ integration (primers 1 and 3), 3’ integration (primers 4 and 2) and wild type DNA template (primers 1 and 2). Primers for parasite18S rRNA were used as loading DNA control. **B)** Comparison of the number of oocysts per midgut and salivary glands (Sg) sporozoites (SPZ) per mosquito between *Pf* NF54^CSP^GFP line and *Pf* NF54 WT parasites (* historical data were used for *Pf* NF54 WT for the comparison). **C)** Percent oocysts expressing GFP under CSP promoter observed in the midguts under fluorescent microscope on day 7 and 11 post *Pf* NF54^CSP^GFP infected blood feed to the mosquitoes. **D)** Western blot analysis of CSP and GFP expression in the lysate prepared from 1 million sporozoites and 10 *Pf* NF54^CSP^GFP infected mosquito midguts of Day 5, 7, 9 and 11 post infected blood feed.

**Supplementary Figure S2:**
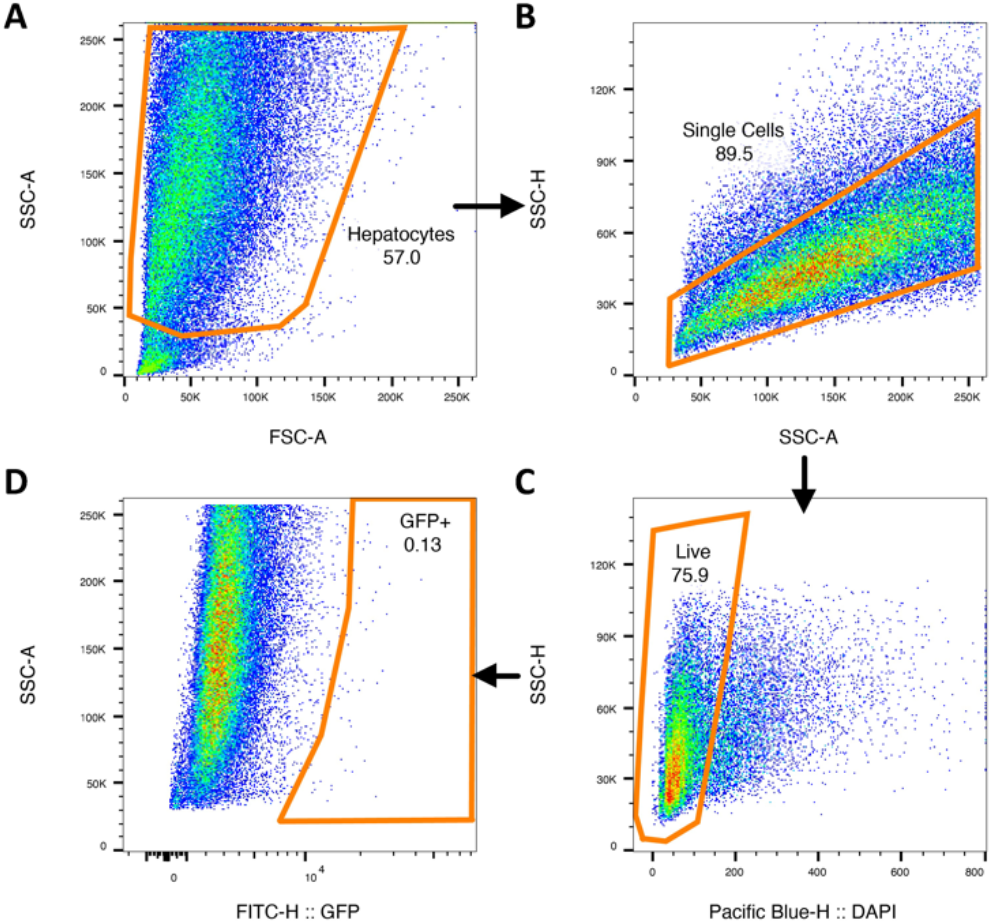
Representative FACS gating for the isolation of Pf NF54^CSP^GFP parasite infected hepatocytes from FRGhuHep mice. **A)** Forward and side scatter gating based on general size to find hepatocytes. **B)** Gating to distinguish single cells from doublets. **C)** Live hepatocytes gating based on DAPI. **D)** Gating of infected primary hepatocytes based on GFP content.

**Supplementary Figure S3:**
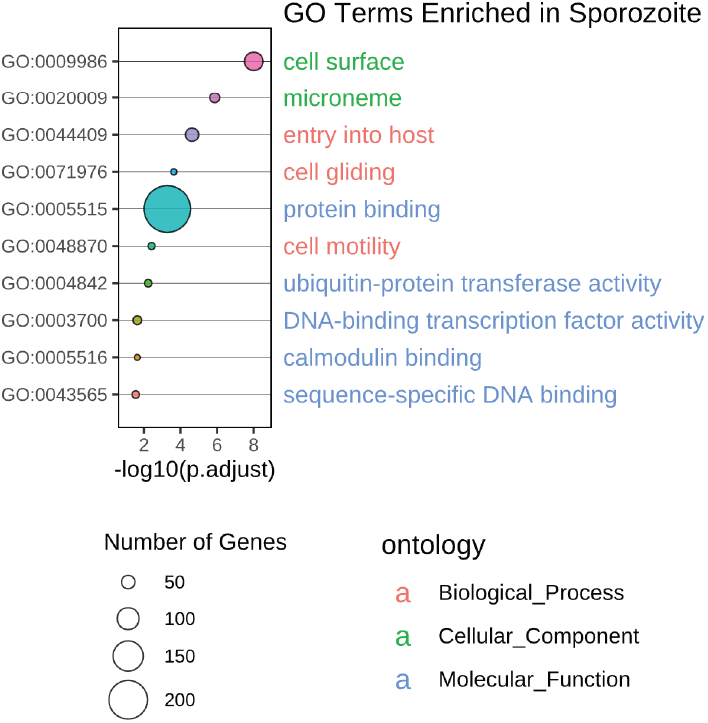
Sporozoites upregulated pathways: Bubble plot showing GOTerm analysis of the upregulated genes in Sporozoites. The size of the circles display by different biological processes is positively correlated with the number of genes involved in each pathway. Pathway plotted have adjusted p-value < 0.05.

**Supplementary Figure S4:**
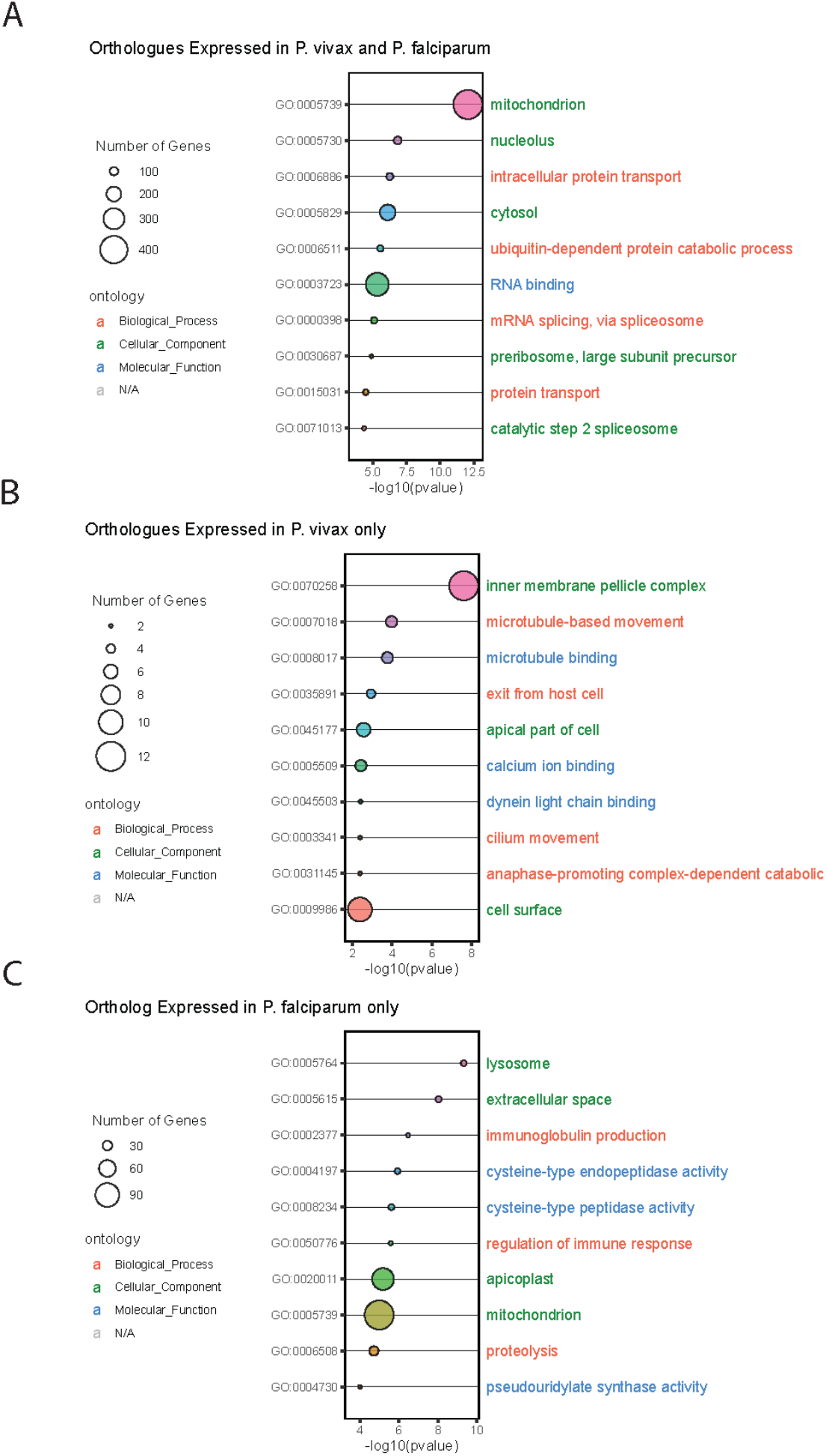
*P. falciparum* and *P. vivax* expressed orthologues and pathways. Bubble plot showing GOTerm analysis of the expressed genes in **A)** *Pf* and *Pv,* **B)** genes expressed only in *Pv* LS transcriptome and **C)** genes expressed only in *Pf*. The size of the circles display by different biological processes is positively correlated with the number of genes involved in each pathway. Pathway plotted have adjusted p-value < 0.05.

